# Genomic rearrangements have consequences for introgression breeding as revealed by genome assemblies of wild and cultivated lentil species

**DOI:** 10.1101/2021.07.23.453237

**Authors:** Larissa Ramsay, Chu Shin Koh, Sateesh Kagale, Dongying Gao, Sukhjiwan Kaur, Teketel Haile, Tadesse S. Gela, Li-An Chen, Zhe Cao, David J. Konkin, Helena Toegelová, Jaroslav Doležel, Benjamin D. Rosen, Robert Stonehouse, Jodi L. Humann, Dorrie Main, Clarice J. Coyne, Rebecca J. McGee, Douglas R. Cook, R. Varma Penmetsa, Albert Vandenberg, Crystal Chan, Sabine Banniza, David Edwards, Philipp E. Bayer, Jacqueline Batley, Sripada M. Udupa, Kirstin E. Bett

**Affiliations:** Department of Plant Sciences, University of Saskatchewan, Saskatoon, Canada; National Research Council of Canada Saskatoon, Canada; USDA-ARS, Small Grains and Potato Germplasm Research Unit, Aberdeen, ID, USA; Agriculture Victoria, Department of Jobs, Precincts and Regions, AgriBio, Centre for AgriBioscience, Bundoora, Victoria, Australia; Institute of Experimental Botany of the Czech Academy of Sciences, Center of the Region Haná for Biotechnological and Agricultural Research, Olomouc, Czech Republic; USDA, ARS, Animal Genomics and Improvement Laboratory, Beltsville, MD, USA.; Department of Horticulture, Washington State University, Pullman WA, USA.; USDA, ARS, Plant Germplasm Introduction and Testing Research, Pullman, WA, USA; USDA, ARS, Grain Legume Genetics and Physiology Research Unit, Pullman, WA; Department of Plant Pathology, University of California Davis, Davis CA, USA; School of Biological Sciences and School of Agriculture, University of Western Australia, Crawley, WA, Australia; International Center for Agricultural Research in the Dry Areas (ICARDA), Rabat, Morocco

## Abstract

Understanding the genomic relationship between wild and cultivated genomes would facilitate access to the untapped variability found in crop wild relatives. We developed genome assemblies of a cultivated lentil (*Lens culinaris*) as well as a wild relative (*L. ervoides*). Comparative analyses revealed large-scale structural rearrangements and additional repetitive DNA in the cultivated genome, resulting in regions of reduced recombination, segregation distortion and permanent heterozygosity in the offspring of a cross between the two species. These novel findings provide plant breeders with better insight into how best to approach accessing the novel variability available in wild relatives.

---

Lentil (*Lens culinaris* Medik.) has been an important grain legume since the dawn of agriculture. Originating in the Middle East and Central Asia, lentil is now grown in many areas of the world as an important source of nutrition domestically or for export. It is rich in fiber, protein, micronutrients, and complex carbohydrates, low in fat and has a low glycemic index. Lentil is a dietary staple in many Middle Eastern and South Asian countries and is gaining popularity in other regions because it is easy to cook and suitable in diabetic, gluten-free, and heart-smart diets. Typically grown in a cereal-based cropping system, lentil improves soil structure and biological diversity, and enhances soil fertility through its nitrogen fixing ability, making it a key crop for sustainable agriculture. Besides its nutritional and environmental benefits, lentil is a valuable cash crop for many farmers around the world. Global lentil production has increased more than five-fold over the past five decades (www.fao.org/faostat), and this upward trend is expected to continue.

With the growing demand for lentil worldwide, genetic improvement for higher and stable yield, improved tolerance to biotic and abiotic stresses, and enhanced nutritional and culinary quality become more critical. Conventional plant breeding strategies have been successful but are resource-intensive, time-consuming and lack precision, especially when it comes to traits that are governed by complex networks of genes. Genomic strategies offer the potential to track and select desirable traits precisely and efficiently accelerate varietal development. Availability of a reference genome sequence enables breeders to make informed selections at earlier stages of the breeding cycle through the use of markers and/or genomic estimated breeding values (GEBVs), thereby accelerating the rate of genetic gain. An initial lentil genome assembly using paired short reads resulted in only 2.8 Gb being assembled and was highly fragmented (Bett, unpublished data). The application of long-read sequencing technology has since supported the construction of a highly contiguous assembly enabling detailed genomic comparisons.

Crop wild relatives are an important source of genetic variation in crop breeding programs and have evolved mechanisms to cope with a wide range of biotic and abiotic stresses. Many of the alleles contributing to the success of these wild plants were lost during domestication, but breeders are often reluctant to cross with wild relatives as the cost and perceived downside of introducing deleterious alleles may outweigh the advantage of the desired allele. With a better understanding of the genomes of crop wild relatives, it should be possible to strategically access novel positive alleles and limit the impact of negative ones.

Genomic structural variations, including copy number variants, presence/absence variants, inversions, and larger segmental duplications, deletions and translocations are present in germplasm collections. Structural variants have long been thought to be important factors in speciation due to their capacity to cause mating barriers that can result in locally adapted gene complexes^1^. Increasingly it is clear that the amount of variation and heritability explained by structural variation is substantial^2, 3^ and plays an important role in breeding efforts involving the use of more diverse or wild germplasm^4, 5^. Even if interspecific crosses result in fertile offspring, genes located within inverted or translocated genomic regions in one parent relative to the other may experience reduced recombination that can extend beyond the rearrangement due to the physical nature of the 3D structures formed by chromatids attempting to pair during meiosis. This leads to linkage blocks of genes that are introduced from the wild parent. These blocks represent linkage drag of unwanted genetic material that can be difficult to eliminate.

Wild lentils have been used in breeding programs for many years as a source of new variability in the cultivated gene pool. Efforts have been made to introgress novel alleles for important traits such as disease resistance, environmental stress tolerance, agronomic and quality traits, from the wild gene pool into elite backgrounds. *Lens ervoides* (Bring.) Grande was initially used as a source of disease resistance, but it also offers variability for seed quality and yield traits^6–8^. While crossing between these two species is possible, large-scale structural differences exist between the two genomes, leading to local uneven pairing and recombination^9^.

Here we present the reference cultivated lentil genome assembly based on CDC Redberry, a Canadian red lentil cultivar^10^ and the reference assembly of IG 72815, a wild *L. ervoides* accession used as a source of genetic variability in breeding programs (Fig. 1). We used these assemblies and an interspecific mapping population to better understand the genomic challenges of accessing genetic variability from a wild relative.

**Figure 1.**
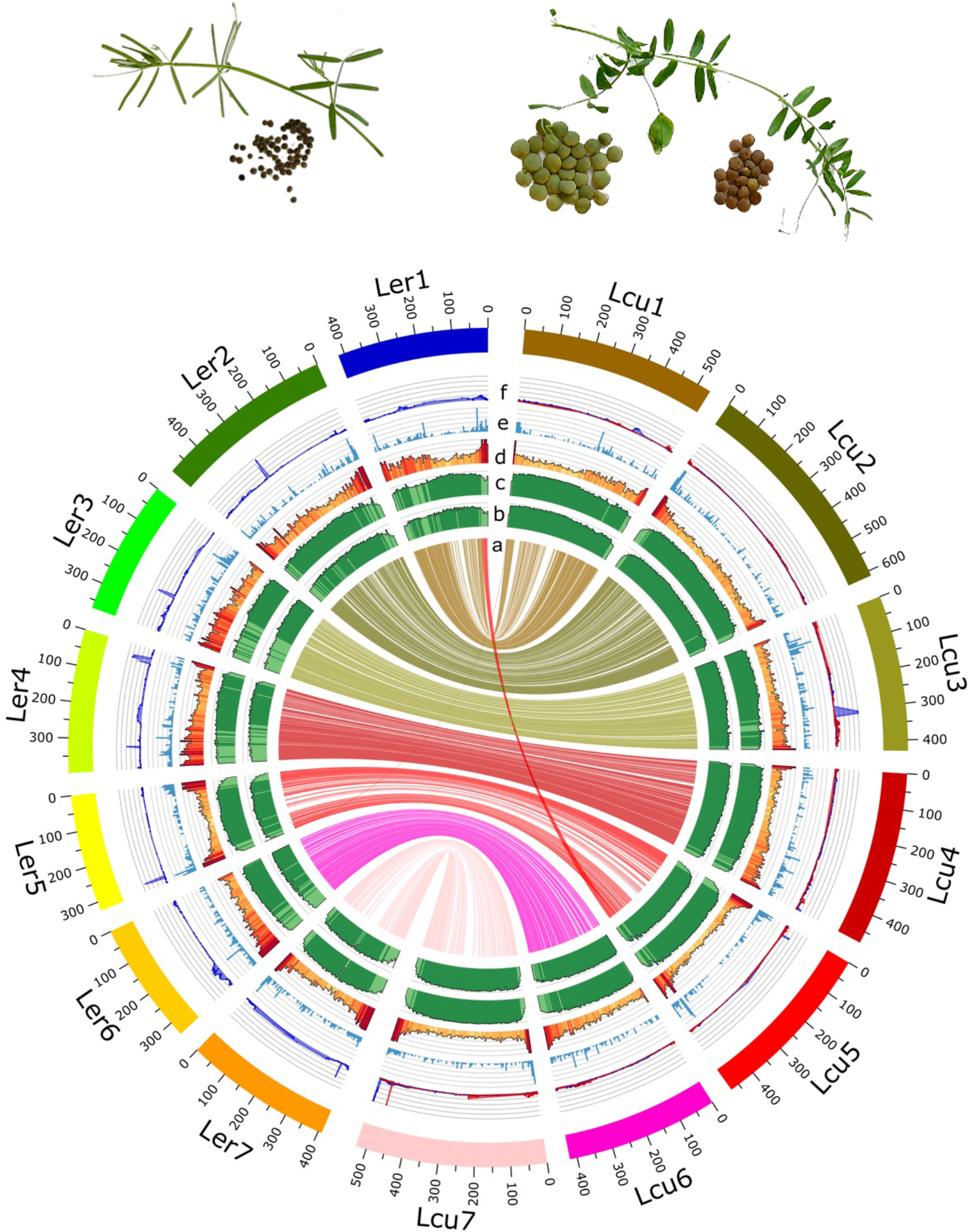
Top =. *Lens ervoides* (left) and *L. culinaris* (right) leaves and seeds. Bottom – genomic characteristics of *L. culinaris* assembly Lcu.2RBY and *L. ervoides* assembly Ler.1DRT. a) syntenic blocks between Lcu.2RBY and Ler.1DRT; b) CG methylation percentage (min. 80%, max.100%); c) proportion of transposable elements; d) gene density (min. 0, max. 200), e) resistance gene analogs distribution (min. 0, max. 20); f) QTL LOD scores for reaction to *Colletotrichum lentis* Race 0 (blue) from *L. ervoides mapping* population LR-66 (mapped to the Ler.1DRT chromosomes, 0<LOD<10) and from the LR-26 interspecific mapping population (mapped to the Lcu.2RBY chromosomes, 0<LOD<6). QTL LOD scores for dehiscence (red) in LR-26.

## Results

### Cultivated and wild genome assemblies

The *L. culinaris* reference genome (Lcu.2RBY; CDC Redberry) was assembled from 54x long-reads, polished using both long reads and additional short reads. Hi-C proximity by ligation and a single genetic map were used to scaffolded contigs into seven pseudomolecules representing 92.8 % of the assembly (ST1). The total assembly covers 3.76 Gbp of the 3.92 Gbp size estimated using K-mer distribution analysis (SF1) and similar to the 4.06 Gbp obtained from flow-cytometry for this species^11^.

The *L. ervoides* reference genome (Ler.1DRT; IG 72815) was assembled from 52x long reads, polished with the long reads and additional short reads. Hi-C proximity by ligation and a single genetic map were used to scaffolded contigs into seven pseudomolecules representing 96.1 % of the assembly (ST1). This assembly is smaller - 2.9 Gbp (ST1), reflecting the smaller size of the genome of this species as predicted by K-mer analysis (∼2.9 Gbp; SF1).

Annotation of the two genomes indicates similar numbers of high confidence genes (ST1) – 39,778 in Lcu.2RBY and 37,045 in Ler.1DRT, representing 94 % and 95 % complete BUSCOs, respectively (ST1). These numbers are similar to those found in pea (*Pisum sativum* L.) ^12^, another sequenced cool season legume with a similar genome size (3.9 Gb, 44,756 complete gene models), and slightly more than what was reported in older legume genome assemblies. There is residual evidence of the whole genome duplication (WGD) common to all papilionoid legumes^13^ that happened ∼56.5 million years ago (Mya). Outside of these few genes, we see little evidence of large-scale segmental duplication in either *Lens* genome. The two lentil species contain similar numbers of disease resistance gene analogues (RGAs), about half the number found in Medicago (*Medicago truncatula* Gaertn.) and slightly more than found in pea (ST2). These genes are located on all chromosomes but are more heavily concentrated on chromosome 3 of both *Lens* species (Fig. 1e, ST2A & ST2B).

### Both cultivated and wild lentil genomes are bursting with repeats

Repeats constitute 82.6 % of the assembled CDC Redberry genome: 3.2 Gb (ST3) out of the 3.9 Gb assembly. In IG 72815, the proportion is lower (78.3 %), with 947 Mb fewer LTR transposons contributing to the smaller size of this genome (ST3). These proportions are much higher than for other diploid legume assemblies such as Medicago^14^ (20-23 %), chickpea (*Cicer arietinum* L.) ^15^ (49 %), pigeonpea (*Cajanus cajan* (L.) Huth.)^16^ (52 %), and common bean (*Phaseolus vulgaris* L.)^17^ (45 %); but in line with what was seen in the 3.9 Gb pea genome assembly^12^ (83 %).

The repetitive regions consist primarily of Class I transposable elements (Lcu.2RBY: 83.6 %; Ler.1DRT:76.7 %) mostly LTRs of the *Ty3/gypsy* (Lcu.2RBY: 63.8 %; Ler.1DRT: 54.3 %) and *Ty1/copia* (Lcu.2RBY: 15.4 %; Ler.1DRT: 16.8 %) types. The 1 Gb difference in the size of these two genomes can be attributed largely to the presence of additional *Ty3/gypsy* elements (Fig. 2, ST3). Most of the gypsy elements are *Ogre* or Tekay and the *copia* types are of the SIRE lineage. *Ogre* elements are large (up to 25 kb) and abundant in legumes, especially those of the Vicieae tribe^18^. These repeats constitute a large portion of the missing 1.1 Gb in our original, predominantly short-read, CDC Redberry genome assembly (https://knowpulse.usask.ca/genome-assembly/Lc1.2).

**Figure 2.**
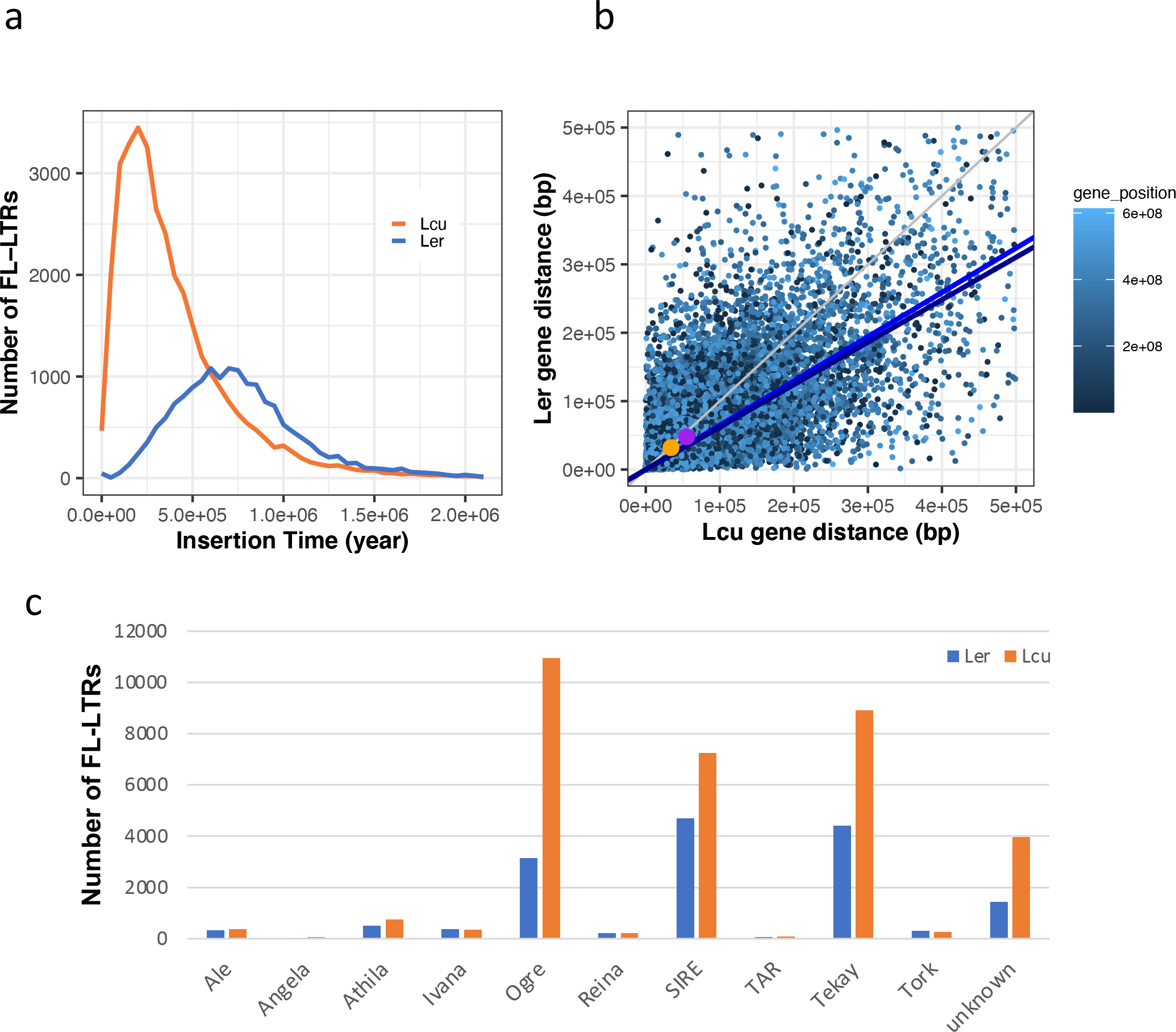
a) Full length LTR-RTs insertion time estimation; b) Genomic distance between two adjacent syntenic genes in Ler.1DRT versus Lcu.2RBY. Regression lines (blue, dark blue) are fitted for whole-genome and pericentromeric region (>100Mb from chromosome-ends), respectively. Orange and purple markers shows the median values of Ler-versus-Lcu gene distance in whole-genome and pericentromeric genes, respectively; c) Classification of full length LTRs.

An analysis of the TE insertion time in each of the genomes revealed that *L. culinaris* had a recent burst of activity – peaking around 0.2 Mya; whereas the peak insertion time for *L. ervoides* TEs was 0.75 Mya (Fig. 2). This is much more recent than is generally observed for cereal crops where the average insertion time for repeats is much more ancient - e.g., barley at 1.4-2.4 Mya^19^ and all wheat sub-genomes at 0.6 (D genome) to 1.5 (A and B genomes) Mya^20^. The younger insertions have not had time to diverge, making these genomes more complicated to sequence, and when combined with the fact that *Ogre* is too large to span with short-read technology, rendering them difficult to assemble without long reads.

Oxford Nanopore read signals were analyzed to assess DNA methylation levels in CpG context. A high proportion of 5-methylated cytosines, 98 % (total 45,160,911 CpG sites) and 95 % (total 30,257,861 CpG sites) were observed in Lcu.2RBY and Ler.1DRT, respectively (ST4). The percentage of methylated cytosines are similar in *Lens*, but much higher compared to soybean (*Glycine* max (L.) Merr.; 51 %) and mungbean (*Vigna radiata* (L.) R. Wilczek var. radiata; 59 %)^21, 22^. The average CpG methylation is 65 % and 62 % in genes and 78 % and 77 % in 1Kb flanking regions for Lcu.2RBY and Ler.1DRT, respectively.

### Phylogeny, mutation rate and divergence from other legumes

The gene coding sequences from 12,963 orthologous genes across *Lens culinaris*, *Glycine max*, *Pisum sativum*, *Medicago truncatula*, *Cicer arietinum* and *Phaseolus vulgaris*, were used to define evolutionary relationships among these leguminous species (Fig. 3). The mutation rate was calculated based on the mean synonymous substitution value of the shared WGD event that occurred ∼56.5 Mya^23^. The results suggest that both *L. culinaris* and *L. ervoides* are evolving at the same rate, and slightly more slowly than *P. sativum* (0.93x) (ST5). However, the two *Lens* species are evolving more rapidly than the legume species with smaller genomes. Despite both *Lens* species having experienced a similar mutation rate, the distance between syntenic pairs of genes is larger in *L. culinaris* than in *L. ervoides* (34.1 Kb vs 32.0 Kb; Fig. 2b), especially in the pericentromeric regions where the difference is 55.4 Kb in *L. culinaris* vs 47.7 Kb in *L. ervoides*. The difference lies in the intergenic repeat proliferation which stretches the physical distance between homologous genes (Fig. 2).

**Figure 3.**
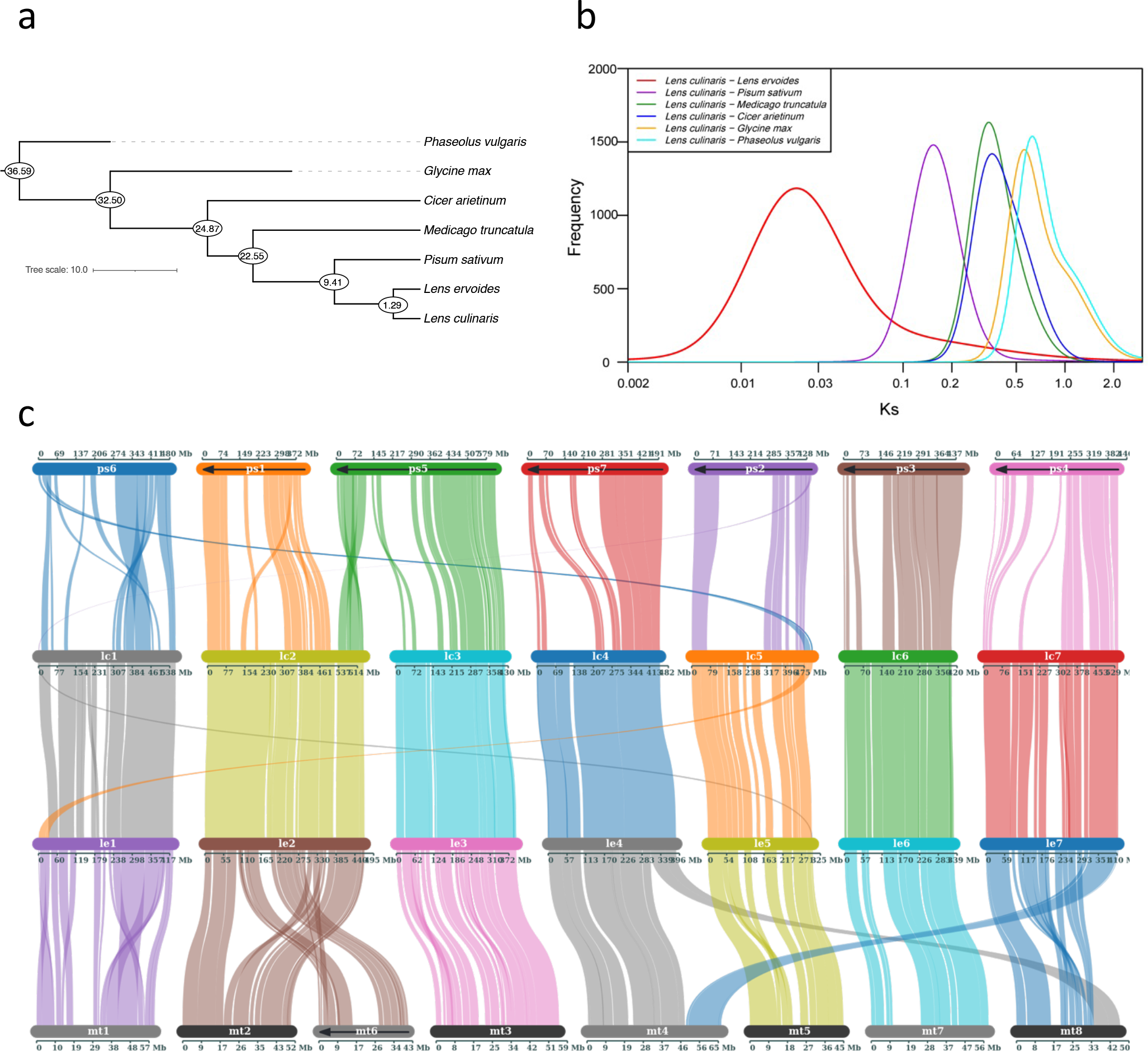
a) Phylogenetic relationship between different legume species; b) Divergence time estimation based on Ks distributions. Mixed model analysis was performed to identify major peaks in Ks values between orthologous pairs; c) Synteny blocks between *Pisum sativum* (Ps), *Lens culinaris* (Lc), *Lens ervoides* (Le), and *Medicago truncatula*(Mt).

Based on a calibrated synonymous substitution rate of 8.3 x 10^-9^ and the geometric mean of the peak observed in each Ks distribution, the ages of divergence between *L. culinaris* and these other legume species were estimated (Fig. 3; ST6; SF2). Lentil diverged from pea and Medicago over 9 Mya and 22 Mya, respectively, and the two *Lens* species diverged almost 1.3 Mya.

### Collinearity with other legumes

Large scale collinearity exists among pea, Medicago, and the two lentil species examined, with several notable translocations and inversions (Fig. 3). A large, inverted, reciprocal translocation exists between chromosomes 2 and 3 in both *Lens* spp. relative to pea chromosomes 1 and 5 but not in Medicago, suggesting this is a feature of the pea genome. The Mt4-Mt8 translocation that is known to exist in the A17 line used for the Medicago genome^24^ is clearly visible relative to *Lens* chromosomes 4 and 7. The middle of Mt8 is inverted relative to the two *Lens* genomes. The extra chromosome in Medicago, Mt6, corresponds to the middle of chromosome 2 for both *Lens* spp. There is a large inversion within chromosome 1 of the two lentil genomes relative to both pea and Medicago.

As noted previously based on genetic linkage maps^9, 25, 26^, *L. culinaris* chromosomes 1 and 5 have an unbalanced, reciprocal translocation relative to both Medicago and pea, but also relative to *L. ervoides*, indicating that the translocation occurred after *L. culinaris* and *L. ervoides* diverged. The assemblies show that not only is there a translocation, but that the fragments are inverted in *L. culinaris* relative to *L. ervoides*. In addition to the major translocation, there are multiple intrachromosomal inversions, and a translocation within each of chromosomes 1 and 7, and an additional inversion on chromosome 5 in *L. culinaris* relative to *L. ervoides* (Fig. 4). There are two smaller inversions, on chromosomes 2 and 6 respectively, but none of notable size in the other chromosomes. With these assemblies we can explore the consequences of these breaks in collinearity more closely.

**Figure 4.**
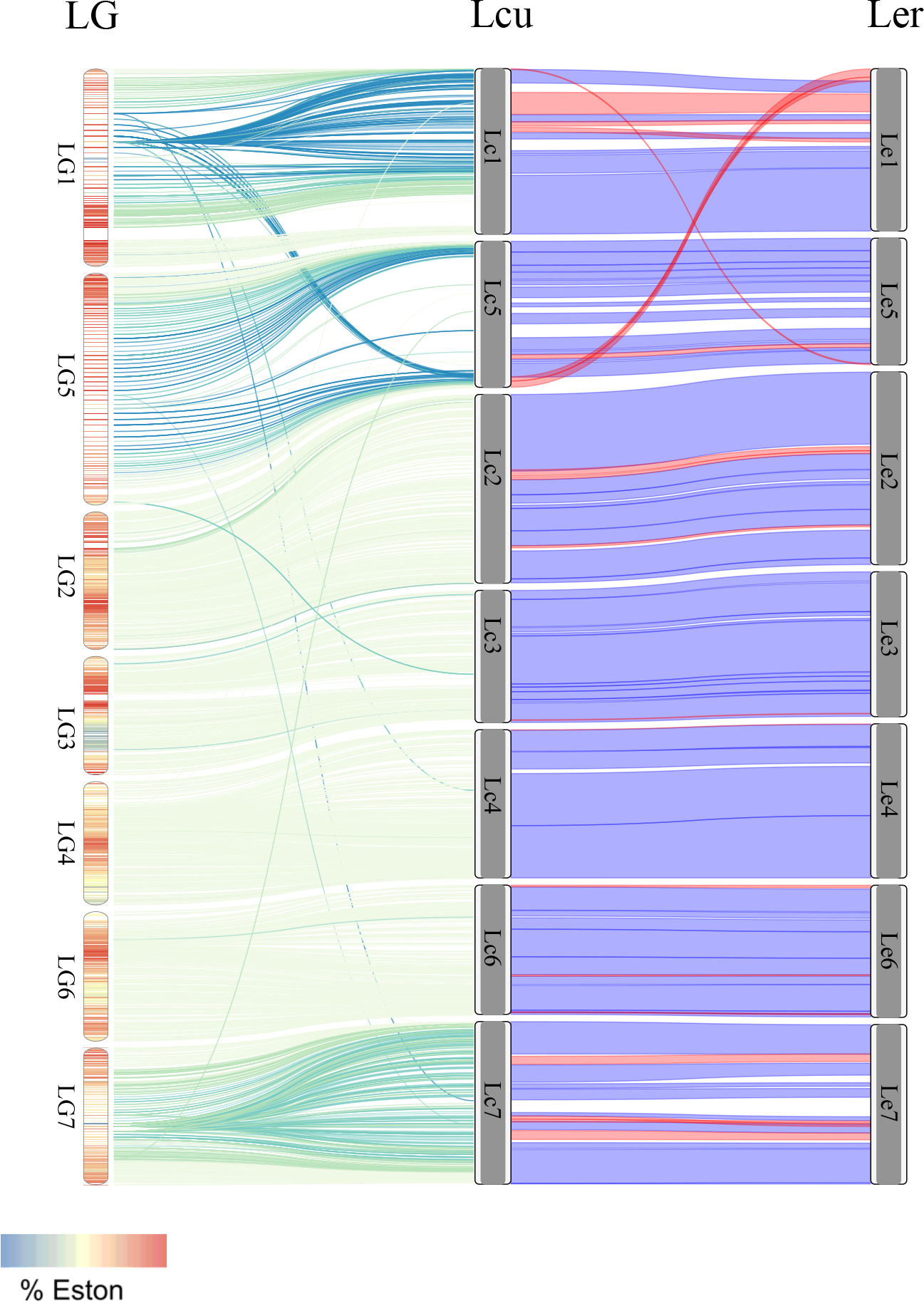
Synteny between LR-26 genetic linkage map (left), L. **culinaris** (Lcu.2RBY) **genome (middle) and L ervoides** (Ler.1DRT) **genome.** LR-26 linkage groups are shaded according to the level of distortion. Lines between the LR-26 linkage groups and Lcu.2RBY are shaded according to the level of heterozygosity observed in LR-26 (green:low,blue:high). Lines between the 2 physical maps are coloured based on conserved synteny (blue: plus strand, red: minus/inverted).

### Breaks in collinearity between *L. culinaris* and *L. ervoides* have consequences for introgression

LR-26 is a recombinant inbred line (RIL) population derived from a cross between *L. culinaris* cv. ‘Eston’ and *L. ervoides* accession IG 72815^7^. This population was created to better understand the genetics of disease resistance in *L. ervoides* but can also be used to examine the consequences of crossing between species with large genomic rearrangements. The 5,491 SNP markers used to create a genetic linkage map of LR-26 were obtained by mapping GBS reads onto the Lcu2.RBY genome and calling SNPs relative to that assembly. The resulting genetic linkage map consists of seven linkage groups: five corresponding to *L. culinaris* chromosomes 2, 3, 4, 6 and 7 and one corresponding to much of chromosome 5 (ST7). The linkage group that corresponds to Lcu.2RBY chromosome 1 contains a region with limited recombination, including a block of 28 binned markers adjacent to a block of another 222 binned markers that contain SNPs that correspond to locations on both chromosomes 1 and 5 of *L. culinaris* (Fig. 4). This region of the map involves pseudo-linkage of the 436-439 Mb region from LcuChr5 mapping between markers within 2-5 Mb of LcuChr1. This region corresponds to where there are large structural variations in one species relative to the other, including the larger 1-5 translocation at one end (Fig. 4). This translocation would likely cause the two pairs of chromosomes to come together during meiosis and, when coupled with the inverted nature of the translocation as well as the other inversions found on those chromosomes, would lead to a reduction in pairing and recombination as well as pseudo-linkage. Large inversions can lead to non-linear pairing which, combined with recombination outside of the loop, will lead to balanced gametes. Recombination within the loop, however, will lead to unbalanced gametes, reduced fertility and a lack of recombinants in that region in resulting offspring. In addition, asynapses will drive the reduced recombination past the breakpoints extending the region of reduced recombination even further. The consequence is that long stretches of binned markers occur in the genetic linkage map and increased recombination occurs outside the affected region, as shown in the linkage groups that correspond to chromosomes 1 and 5 (Fig. 4). A further impediment to pairing and recombination would be the larger physical distances between genes in *L. culinaris* (Fig. 3), which would lead to additional complications in pairing, again limiting opportunities for recombination.

After generating the genetic linkage map of LR-26, the distorted markers that had been removed were added back based on their location in the reference genome to determine if there was segregation bias towards one or the other parental genome (Fig. 4). Overall, in regions where distortion exists, it is towards the *L. culinaris* parental allele. The only exception is a region on chromosome 3 from 345-384 Mb where there is a slight bias towards *L. ervoides* alleles. This region corresponds to a region of the intraspecific *L. ervoides* genetic linkage map, LR-66, which contains an inversion relative to the Ler.1DRT assembly (SF3), suggesting that this region may be susceptible to rearrangements that could lead to altered segregation patterns. Much of chromosomes 1 and 5 are biased towards the *L. culinaris* alleles and in places the distortion was extreme (>0.8 *L. culinaris* allele state). During the development of LR-26, efforts were made to reduce any selection bias towards the domesticated parent including scarifying seed prior to planting to eliminate physical dormancy, collecting seed before pods shattered and not eliminating the lines that took longer than the rest to emerge or to mature. Clearly selection is occurring, if not at the whole plant level, then during meiosis. Bias towards the one parent could be due to centromere drive^27^ which is usually associated with increased numbers of specific repetitive elements in centromeric regions in one parent over the other. There are slightly more lentil-specific centromeric satellite repeats in the *L. ervoides* parent (1.3Mb vs 1.2Mb in *L. culinaris*) (ST8), however. Total centromeric CRM-clade retrotransposon sequence is equal in both species (3.1Mb) as well, suggesting this element is likely not involved.

The location of centromeric satellite repeats associated with centromeric chromatin does not always line up with the expected centromeres. For instance, there are three regions on chromosome 3 of both genomes that contain these elements (ST8; SF3 & 4). Chromosomes 2 and 7 of Ler.1DRT have two regions but there is only one assembled in the corresponding Lcu.2RBY chromosomes. Non-centromeric loci with these satellite repeat sequences have been identified in *Pisum sativum*^28^. There is no signal of centromeric repeats on chromosome 4 of Lcu.2RBY nor on chromosomes 4 or 5 of Ler.1DRT which could be due to incomplete scaffolding of centromeric regions; however, potato research reports also indicate a lack of centromeric repeats on some centromeres^29^.

Plotting the LR-26 markers across the genome in terms of percent of heterozygous calls (Fig. 4), chromosomes 1, 5 and to a lesser extent 7, all have regions with elevated numbers of individuals that have retained a heterozygous state, even after more than six generations of selfing. The regions on chromosomes 1 and 5 also correspond to the regions that are distorted towards the *L. culinaris* allele. In the case of chromosome 1, this involves the first 325Mb of sequence which includes the region with the translocation as well as several inversions in *L. culinaris* relative to *L. ervoides*. Beyond the region with the structural rearrangements, the number of heterozygotes drops to zero. Linkage group 5 corresponds to chromosome 5 but most of the markers are found clustered at the two ends with high levels of recombination between markers. Linkage group 5 has an elevated heterozygous state across most of its length, apart from the region adjacent to the translocated region. Chromosome 7 has multiple inversions in one species relative to the other through the interior portions of its length, which are likely responsible for the increased level of heterozygosity seen in the same region in the genetic linkage map.

The strong linkage and relatively high levels of heterozygosity between the set of markers before the translocation breakpoint on chromosome 5 and the region central to chromosome 1 suggests the presence of permanent translocation heterozygotes within the interspecific RIL population. Heterozygous calls are in the same RILs across the translocation region (ST7). Inversions and large structural rearrangements already present in *L. ervoides* relative to *L. culinaris* contribute to the lack of recombination across the region. Differences in gene content between chromosomes, particularly around the translocation breakpoint, would contribute to maintenance of a structurally heterozygous state. Conversely, linkage group 7, which has lower levels of heterozygosity throughout, has only inversions. One possible scenario could be that genes within a rearrangement that are identical by descent could have differentially sub-functionalized over time, leading to a situation where if both are not present, there will be reduced viability of the offspring. If gene content requirements drive structural heterozygosity, it is possible that the inversions are less likely to impact genes compared to a translocation, explaining the lower levels of heterozygosity in linkage group 7. Alternatively, the inversions on chromosome 7 are relatively younger and any genes have yet to sub-functionalize. It is worth noting that the low levels of recombination observed on linkage group 7 in the intraspecific *L. ervoides* map LR-66 and the intraspecific *L. culinaris* map LR-01 (SF3&4), in addition to the other evidence surrounding the chromosome 1/5 breakpoint, may indicate that there are other rearrangements on chromosome 7 even within intraspecific crosses.

## Discussion

*Lens* is a genus of tribe Vicieae, which includes other diploid genera that tend to have large genomes (*Vicia* – 1.8 – 13.3 Gb; *Pisum* – 4.4-4.9 Gb; *Lathyrus* – 3.4 – 14.6 Gb; https://cvalues.science.kew.org/). Assembling these genomes has proven a challenge due to their repetitive nature. We have now assembled almost the entire cultivated lentil genome. This is a vast improvement over the original short-read cultivated lentil assembly which was missing 32 % due to collapsed repetitive regions that could not be spanned and resolved with short-read technology. We also assembled the smaller genome of its wild relative, *L. ervoides*, as a step to better understanding genome evolution in *Lens* and other legumes, as well as to gaining knowledge of the consequences of trying to access genetic variability through interspecies crossing with crop wild relatives.

These genomes are large for diploid plants, mostly due to the presence of large numbers of repeats. The repeats are also what constitute the difference in genome size between the two lentil species genomes and are likely driving the rearrangements observed between them. Chromosome-level structural variation such as translocations and inversion are considered drivers of genome evolution and speciation. Comparative mapping indicates several rearrangements occurred after the *Lens spp*. diverged from pea and other cool season legumes, but before *L. culinaris* and *L. ervoides* diverged. Other rearrangements are unique to one or the other of these *Lens* genomes (Figs. 1 & 4). While large-scale inter-chromosomal rearrangements are not common, inversions and smaller intrachromosomal translocations are evident on all chromosomes, but are particularly prevalent on chromosomes 1, 5 and 7. These rearrangements have likely contributed to the divergence of these species because they can lead to reproductive isolation if they are large enough to interfere with pairing and recombination during meiosis.

Rearrangements in the cultivated species relative to the wild clearly affect pairing and recombination as well as gamete survival in the interspecific hybrids, and this leads to consequences for potential introgression of genes from the wild species into cultivated lentil. Transferring genes found on *L. ervoides* chromosomes 2, 3, 4 and 6 to cultivated lentil should be relatively successful. However, genes of interest on chromosomes 5 and 7 and the long arm of 1 could be difficult to introgress because of the negative consequences of extensive linkage drag, possibly coupled with permanent heterozygosity.

To assess the potential to introgress traits, we looked at accessing disease resistance genes from *L. ervoides* for breeding disease resistant lentil cultivars. Resistance to *Colletotrichum lentis* race 0, causal agent of anthracnose was first identified in *L. ervoides* but has not been found in *L. culinaris*. The intraspecific RIL population LR-66 (L01-827a x IG 72815) was developed to map resistance genes in *L. ervoides*. Bhadauria et al.^26^ mapped four QTL for resistance within *L. ervoides* using a genetic linkage map of LR-66 generated after mapping GBS reads to version 0.8 of the CDC Redberry genome assembly. Re-mapping those GBS sequences to the current *L. ervoides* assembly (Ler.1DRT) resulted in more mapped markers and a map that aligns with the genome assembly. This allowed us to place the QTL in the context of their native genome and to narrow down the regions containing candidate resistance genes. QTL for resistance from IG 72815 were identified on chromosomes 3, 4 and 7 (Fig. 1) and fall in regions containing multiple RGAs. The QTL on chromosome 4 were not identified in the original analysis^26^ due to a lack of markers in the region, demonstrating the power of using the native genome for read mapping and SNP calling. Additional resistance QTL were mapped to chromosomes 2 and 5 that came from the other parent, L01-827a, as was found in the original analysis.

The interspecific (*L. culinaris* x *L. ervoides*) RIL population LR-26 was screened for resistance to anthracnose (*C. lentis* race 0) and QTL were observed on the two linkage groups that align with chromosomes 3 and 7^30^. These also line up with the two LR-66 intraspecific linkage groups containing QTL (Fig. 1) that came from the common parent, IG 72815. Since chromosome 3 does not contain large-scale structural variation in *L. ervoides* relative to *L. culinaris*, it should be possible to introgress that segment and reduce linkage drag through normal recombination and selection. The distortion towards the *L. culinaris* parent alleles in that region (Fig. 4) would suggest it would be prudent to increase the number of offspring to be able to identify candidate lines for further backcrossing to recover the *L. culinaris* state in surrounding regions. The LR-66 QTL on chromosome 7 of *L. ervoides* appears to be more proximal than the one on linkage group 7 of LR-26 (Fig. 1), but this may be an artefact of aligning an interspecific linkage map to the Lcu2.RBY genome. The LR-26 QTL on chromosome 7 is near the end (517-522 Mb) where there appears to be no inversions and in a region with only mild distortion towards the *L. culinaris* allele (Fig. 4). The elevated level of heterozygosity and reduced recombination along much of the rest of this chromosome, however, will mean screening of a much larger population would be needed to eliminate unwanted *L. ervoides* alleles. For example, a pod dehiscence QTL mapping to the region around 430 Mb on Lcu2.RBY chromosome 7 (Fig. 1) will need to be separated from the resistance locus at the end of this linkage group. The two LR-66 QTL on chromosome 4 of *L. ervoides* do not appear to have an effect in LR-26. Given that there does not appear to be any structural reason for this (Fig. 4), it could be due to an epistatic effect of loci from L01-827a that are not present in the interspecific population. Alternatively, the original source of IG 72815 may be heterogeneous, and the actual parent plant used to create LR-26 did not have the positive alleles at those loci, although this is unlikely to happen at two distinct loci.

Sequencing and assembling whole genomes, even those as large as lentil, is becoming easier and less costly. Having genome assemblies for wild and cultivated lines will make accessing the genetic variability available in wild relatives more precise and economical. Mapping sequencing reads to their own genome rather than relying on alignment to a related domesticated species results not only in more markers, but improved QTL analyses and a better chance of candidate gene identification. Understanding where genes of interest lie in the context of the structural rearrangements of cultivated and wild genomes would guide the choice among strategies for their introgression from wild relative to cultivated ones, either by introgression crosses or from recent advances in technologies such as CRiSPR-Cas9 that have led to the possibility of inverting regions in a genome^31^. If this were implemented in lentil, we could envision accessing genes that are currently inaccessible in the inverted regions of a wild parent relative to the cultivated genome and simultaneously reducing linkage drag.

## Online Methods

### Plant Material

CDC Redberry is a small red lentil (*L. culinaris*) variety bred at the University of Saskatchewan, Canada, and registered for cultivation (CFIA #5771) in 2003^10^. IG 72815 is a wild lentil (*L. ervoides*) line originally obtained from the gene bank at the International Centre for Agricultural Research in the Dry Areas (ICARDA). It has been used in interspecific crosses with *L. culinaris* to transfer resistance to multiple diseases^7^ and is one of the parents of the LR-66 intraspecific RIL population^26^. For both genotypes, a single plant was selfed to create the genetic stock for further genomic studies. Tissue from a single plant was collected and high molecular weight DNA extraction was performed from nuclei^32^ using the salting-out method from 10x Genomics^33^ for all sequencing for both species.

### Genetic Mapping

The data used to generate an *L. culinaris* intraspecific genetic linkage map of the RIL population LR-01, derived from a cross between cv. ‘CDC Robin’ and ILL 1704^34^ were revisited and additional markers were included for anchoring and orientation of the CDC Redberry scaffolds into pseudomolecules. The recombination matrix for the map can be found at https://knowpulse.usask.ca/Geneticmap/2695342.

GBS reads of the *L. ervoides* intraspecific RIL population LR-66^26^, derived from the cross L01-827a x IG 72815, were mapped against the Ler.1DRT contigs and used to organize the contigs into bins for HiC scaffolding. The QTL analysis described in Bhadauria et al.^26^ was re-run using their disease reaction scores for *C. lentis* race 0 and the new linkage map. The recombination matrix for the map can be found at https://knowpulse.usask.ca/Geneticmap/2695193.

LR-26 is a population of F7-derived recombinant inbred lines derived from a cross between a single plant of cv. ‘Eston’ (*L. culinaris*) and a single plant from *L. ervoides* accession IG 72815^7^. The genetic linkage map construction and QTL analysis for reaction to *C. lentis* race 0 are described in Gela et al.^30^ and available at https://knowpulse.usask.ca/Geneticmap/2691115. The recombination matrix for the map used for analyses can be found in ST10. Dehiscence data were from Chen^35^.

### Hi-C

Hi-C reactions were performed according to the Arima Hi-C Kit (PN A510008) user guide for plant tissue. The resulting purified Arima Hi-C proximally ligated DNA was used to prepare Illumina-compatible sequencing libraries. Ligated DNA was first sheared to an average size of 600 bp, fragments from 400-1000 bp were selected using SPRI beads and biotinylated fragments were enriched. Libraries were then prepared using Swift Biosciences Accel-NGS 2S Plus DNA Library Kit (PN 21024) and Swift Biosciences 2S Indexing Kit Set A (PN 26148). The final libraries underwent standard quality control (qPCR and Bioanalyzer) and were sequenced on the Illumina HiSeq 2500 following manufacturer’s instructions using Rapid chemistry. HiC contact map results are displayed in SF5.

### Optical map

High molecular weight (HMW) DNA was prepared from 5.6 x 10^6^ intact mitotic metaphase chromosomes (∼6.6 μg DNA) purified by flow cytometric sorting. Preparation of suspensions of chromosomes from synchronized root tips of young seedling and chromosome sorting were done as described by Vrána et al.^36^ HMW DNA was prepared from the isolated chromosomes following Šimková et al.^37^ and labelled using Nick Label Repair and Stain (NLRS) DNA Labeling Kit (Bionano Genomics, San Diego, USA) at Nt.BspQI sites (GCTCTTC motif) as described by Staňková et al.^38^ The analysis of labelled DNA on the Irys platform (Bionano Genomics) using six Irys chips yielded 1154 Gbp of raw data greater than 150 kbp representing 281-fold genome coverage that was used for *de novo* assembly of the lentil optical genome map.

### Pseudomolecule assembly of reference genomes

Assembly of the Lcu.2RBY contigs was done using smartdenovo (https://github.com/ruanjue/smartdenovo/) with 34x coverage of PacBio SMRT and 20x coverage of Oxford Nanopore Technology (ONT) reads. Contigs were polished with racon^39^, using three rounds of long-read data mapped against them and one round of Illumina short read data (10x coverage). Five lanes of Hi-C data were generated, and a first pass of scaffolding and breaking of chimeric sequence was carried out using SALSA^40^, followed by scaffolding using an optical map using Irys-scaffolding^41^. Scaffolds were assigned to chromosome bins using a genetic map from the LR-01 RIL population exome capture data and ordered and oriented within each bin using ALLHiC^42^. Pseudomolecule assemblies were individually visualized and manual corrections to fix telomere tethering made using Juicebox (https://github.com/aidenlab/Juicebox). Assembly of the Ler.1DRT contigs was done using smartdenovo with an estimated 52x coverage of ONT reads. Contigs were polished with racon^39^, using two rounds of long-read data and one round of Illumina short read data (10x coverage). The GBS-based genetic map for the LR-66 population was used to assign contigs to chromosome bins. Two lanes of Hi-C data were generated and used to order and orient contigs within each bin using ALLHiC^42^, as well as using the software’s rescue mode to assign contigs that contained no markers in the genetic map to bins. The Hi-C data was also passed through SALSA^40^ and used to break several chimeric contigs. Pseudomolecule assemblies were individually visualized and manual corrections to fix telomere tethering issues made using Juicebox.

### Genome size estimates and quality assessments

Illumina short read sequences from *L. ervoides* accession L01-827a^43^ were used for genome size estimation as none were available for IG 72815. Processed Illumina short-read data (∼40X depth) from *L. culinaris* CDC Redberry and *L. ervoides* L01-827a were provided to Jellyfish^44^ v2.2.7 with “-C -m 21 -s 5G –min-quality=25” parameters to generate k-mer (K=21) frequency distribution. The output histograms were analyzed using GenomeScope^45^ to estimate the genome size, heterozygosity level, error rates, and repeat fraction. The presence of conserved plant orthologs was evaluated using BUSCO (v4.0.4) with the Fabales database 10 lineage.

### Gene and RGA Annotation

Strand-specific RNA-Seq data (50.7 Gb) was generated on a MiSeq platform (2x250bp) from seven tissue samples: flowers, leaflets, seedling roots, root nodules, seedlings, etiolated seedlings, and flower buds to assist with the annotation of protein coding genes in *L. culinaris* (ST9). For *L. ervoides*, RNASeq data from *Ascochyta lentis*-inoculated samples^46^ (Sari et al., 2018) were used for annotation. Adapter sequence and low-quality bases were trimmed using Trimmomatic^47^ with the parameters ILLUMINACLIP:2:30:10 LEADING:20 TRAILING:20 SLIDINGWINDOW:4:15 MINLEN:50. The processed data were aligned to the appropriate reference genome using RNAStar^48^ with --alignIntronMax 50000 -- outFilterMismatchNoverReadLmax 0.03 --outFilterMatchNminOverLread 0.95 parameters and subsequently assembled by Trinity software^49^ using reference-guided approach with the -- genome_guided_max_intron 50000 parameter. The assembled transcripts and additional Illumina PBA Blitz lentil transcript data (data associated with *L. culinaris* genome project PRJNA343689 at NCBI), plus sequences from the *M. truncatula* (v4.0) protein database, and the *ab initio* predictors (SNAP^50^ and Augustus^51^, configured in hint-based mode) were used to generate initial gene models. The output from MAKER-P was processed using PASA (v2.3.3)^52^ to further incorporate the transcript alignment evidence into the initial gene annotation. Functional annotation was performed by both BLASTP against the Uniprot-Plant (SwissProt and TrEMBL) and HMMSearch against PFam databases. High confidence genes are defined as a subset of annotated genes with a hit 1e-2 or better to PFam database and/or with combined expression value of TPM>1.

Resistance gene analogs (RGAs) were annotated for both genomes using the RGAugury pipeline^53^ searching against RGA domains in Pfam, Gene3D, SMART, and Superfamily databases with an e-value cutoff = 1e^-5^.

### *Phylogenetic analysis and comparison with Medicago truncatula* and *Pisum sativum*

The gene coding sequences from legume species *L. culinaris*, *L. ervoides*, *Glycine max*, *Pisum sativum*, *Medicago truncatula*, *Cicer arietinum* and *Phaseolus vulgaris*, were compared to construct a supermatrix consisting of 12,679 orthologous genes in a concatenated alignment of 15,886,496 bp, which was used to define evolutionary relationships among these species.

We ran MCScanX^54^ on *L. culinaris*, *L. ervoides*, *Pisum sativum*^12^ and *Medicago truncatula*^55^, requiring a minimum block size of 10 genes. Interchromosomal blocks representing the remainders of the ancestral legume duplication were removed from the plot and the results visualized with Synvisio^56^.

### Repeat analysis

We used a combination of *de novo* and homology-based methods^17, 57^ to annotate transposons in the *L culinaris* genome. To identify LTR retrotransposons, the genome sequence was analyzed with LTR-Finder^58^ using default parameters, but we set 30-bp for minimum LTR length and minimum distance between the LTRs, and 5,000-bp for maximum LTR length. All output sequences were manually inspected to determine the exact boundaries of the characterized LTR retrotransposons and to exclude incorrectly annotated sequences such as tandem repeats. The annotated LTR retrotransposons were used for BLASTX searches to determine their superfamilies. The terminal repeat retrotransposons in miniature (TRIMs) were defined by our previous criteria: 1) element sizes were less than 1,500 bp; 2) at least two complete copies flanked different target site duplications (TSDs) in the genome; and 3) the retroelements encoded no proteins^59^. To annotate long interspersed nuclear elements (LINEs), the protein sequences of published LINEs (http://www.girinst.org/repbase) were used to search against the Lcu.2RBY genome, all significant hits (E value < 1 X e^-15^) and the 10-kb flanking sequences (5-kb on each side) for each hit were extracted and manually examined for polyA tails and TSDs. The short interspersed nuclear elements (SINEs) were annotated by the SINE-Finder^60^ with default settings, and the output sequences were checked for polyA and TSDs. For DNA transposons, the conserved domains for transposases from different DNA transposon superfamilies were used to conduct TBLASTN searches against the lentil genome. The target and flanking sequence (5-kb on each side) were extracted from the genome, the complete DNA transposons were determined by terminal inverted repeats (TIRs) and TSDs. Furthermore, MITEs-Hunter software^61^ was also used to identify small DNA elements that encoded no transposase protein.

All annotated transposons were combined and used as a transposon library to screen the *L. culinaris* and *L. ervoides* genomes using RepeatMasker (http://www.repeatmasker.org) with default settings except that we used the nolow option to avoid masking low-complexity DNA or simple repeats. Transposons were summarized based on names, subclasses and classes, and overlapping regions in the RepeatMasker output file were counted once, and the transposon proportions were calculated based on the coverage of different transposons and gap sequences were excluded for the total genome size.

LTRharvest^62^ and LTR_FINDER^58^ were used to identify LTR retrotransponsons (LTR-RTs) in *L. culinaris* and *L. ervoides* genomes, respectively, using default parameters. The outputs from the software were further analyzed using LTR_retriever^63^ to provide annotation of full-length LTR-RTs. The annotated TE sequences were classified using TEsorter^42^ to provide clade-level classification by searching against the REXdb^64^ database. TE insertion dates were predicted using LTR_retriever using a substitution rate equal to twice the mean synonymous substitution value calculated for each species as proposed by Barghini et al.^65^ and Ma and Bennetzen^66^.

### CpG Methylation

ONT raw reads of *L. culinaris* and *L. ervoides* were aligned to the corresponding genome sequences using minimap2 (-x map-ont parameter). Nanopolish^67^ was used to call methylated (5mC) and unmethylated bases in the CpG context.

### Centromeric sequence analysis

Sequences for both lentil-specific satellite sequences associated with CENH3 chromatin^28^ and a broader selection of CRM-clade chromovirus LTR sequences associated with centromeric satellite regions from *Medicago* and *Pisum sativum*^68^ were used in a BLASTN search of both genomes. Hits scoring an e-value less than 1 X e^-10^ were extracted for estimation of centromere location.

**Table 1.** Genome assembly and annotation statistics of *L. culinaris* and *L. ervoides*.

|  | *L. culinaris* | *L. ervoides* |
| --- | --- | --- |
| Total assembly length | 3.76 Gbp | 2.87 Gbp |
| Total contig number | 6,094 | 2,291 |
| Contig N50 (L50) | 1.3 Mb (841) | 4.7 Mb (176) |
| Total Scaffold number | 2,075 | 1,141 |
| Scaffold N50 (L50) | 482 Mb (4) | 396 Mb (4) |
| Percent in pseudomolecules | 92.81% | 96.14% |
| Number of annotated genes | 58,243 | 60,296 |
| Number of high confidence genes | 39,778 | 37,045 |
| Transposable element coverage | 3.20 Gbp | 2.24 Gbp |
| Percent transposable element | 85.0% | 78.3% |
| Complete BUSCOs | 94.4% | 95.1% |

## Author contributions

KEB, LR, CC (Canada) and AV conceived the study and managed the project.

LR and CSK assembled and annotated the genomes, did the comparative analyses and carried out bioinformatic analyses. SK (Canada) carried out the mutation rate and phylogenomic analyses, DG carried out the repeat analyses, ZC and BR carried out the RGA analyses.

AV, SU, KEB and SK (Australia) contributed genetic resources, DM, JH, CC (USA) and RM contributed PacBio sequencing data, DC, SB and SK (Australia) contributed short read sequencing and transcript data, DK contributed Hi-C and ONT sequencing data, HT and JD carried out the flow sorting of chromosomes and Bionano optical mapping. TH, L-AC, TSG, RS, SB and AV contributed genetic linkage maps, QTL analyses and phenotypic data. DE, PB, JB contributed isolated chromosome and population sequence data.

KEB, LR, CSK, SK (Canada), and DG co-wrote the first draft of the manuscript. CC (USA), RM, AV, DE and VP edited the manuscript.

## Acknowledgements

We thank Christine Sidebottom, Yasmina Bekkaoui, Akiko Tomita, Abdullah Taher, Shimna Sudheesh and Tracie Webster for DNA/RNA preparations, library preparations for sequencing and HiC analysis. Also, Jan Vrána, Zdeňka Dubská, Romana Šperková and Jitka Weiserová for the preparation of chromosome suspensions and flow cytometric sorting.

Financial support from the Saskatchewan Pulse Growers (USask group), Northern Pulse Growers Association, USA Dry Pea & Lentil Council (WSU/USDA group), Grains Research and Development Corporation and Agriculture Victoria (AgVic group), Australian Research Council LP160100030, DP160104497(UWA group), CGIAR Research Program on Grain Legumes (ICARDA/UCDavis group). HT and JD were supported by ERDF project ‘Plants as a tool for sustainable global development’ (No. CZ.02.1.01/0.0/0.0/16_019/0000827). SK (Canada) and DK were supported from the National Research Council Canada through the Sustainable Protein Production Program.

## Supplemental Files

**Supp Fig 1.**
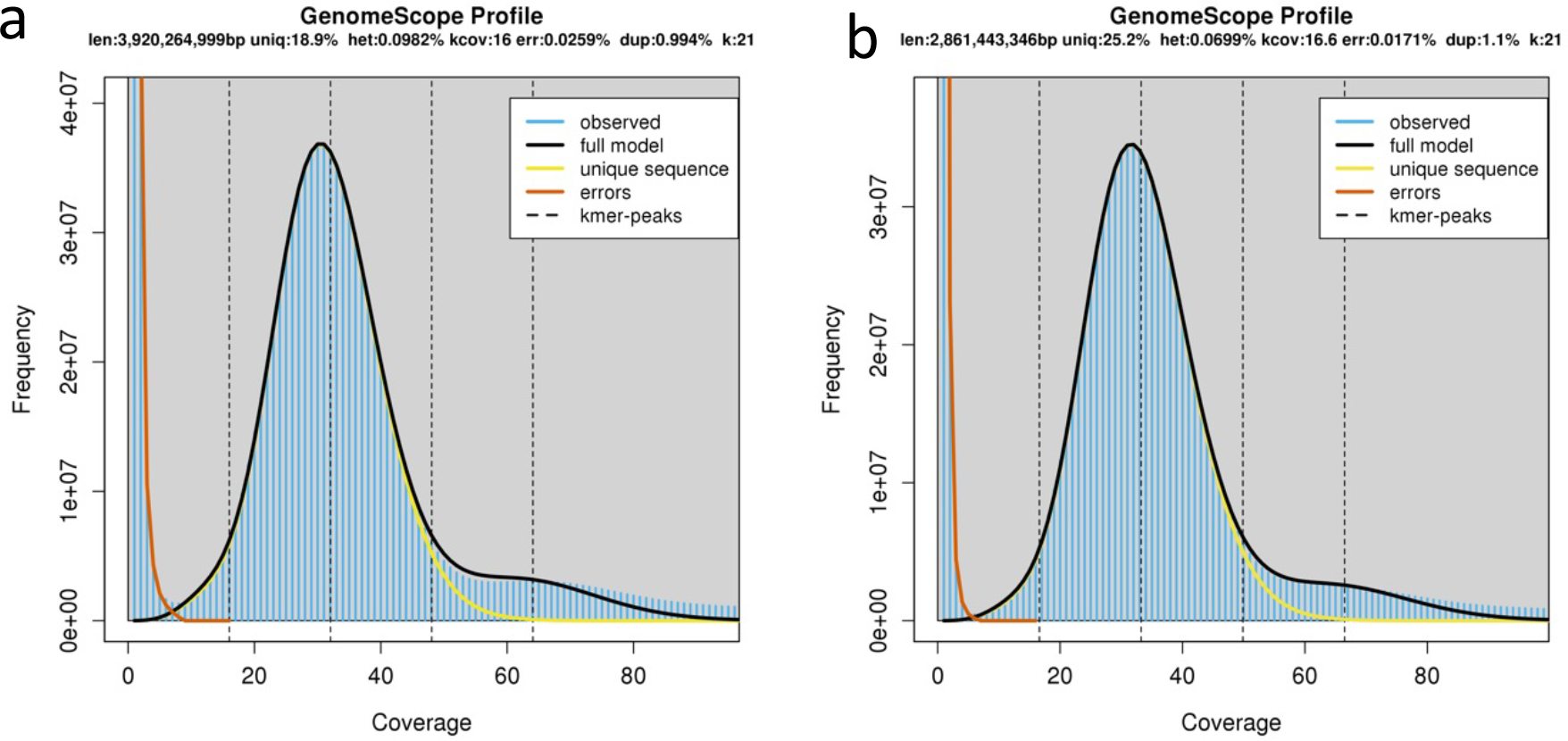
Genome size prediction (K=21) for a) Lcu.2RBY and b) Ler.1DRT using unassembled reads. Reads used for Ler.1DTR were from a different line, L01-827a, due to a lack of sufficient short reads from IG 72815 to do the analysis.

**Supp Fig 2.**
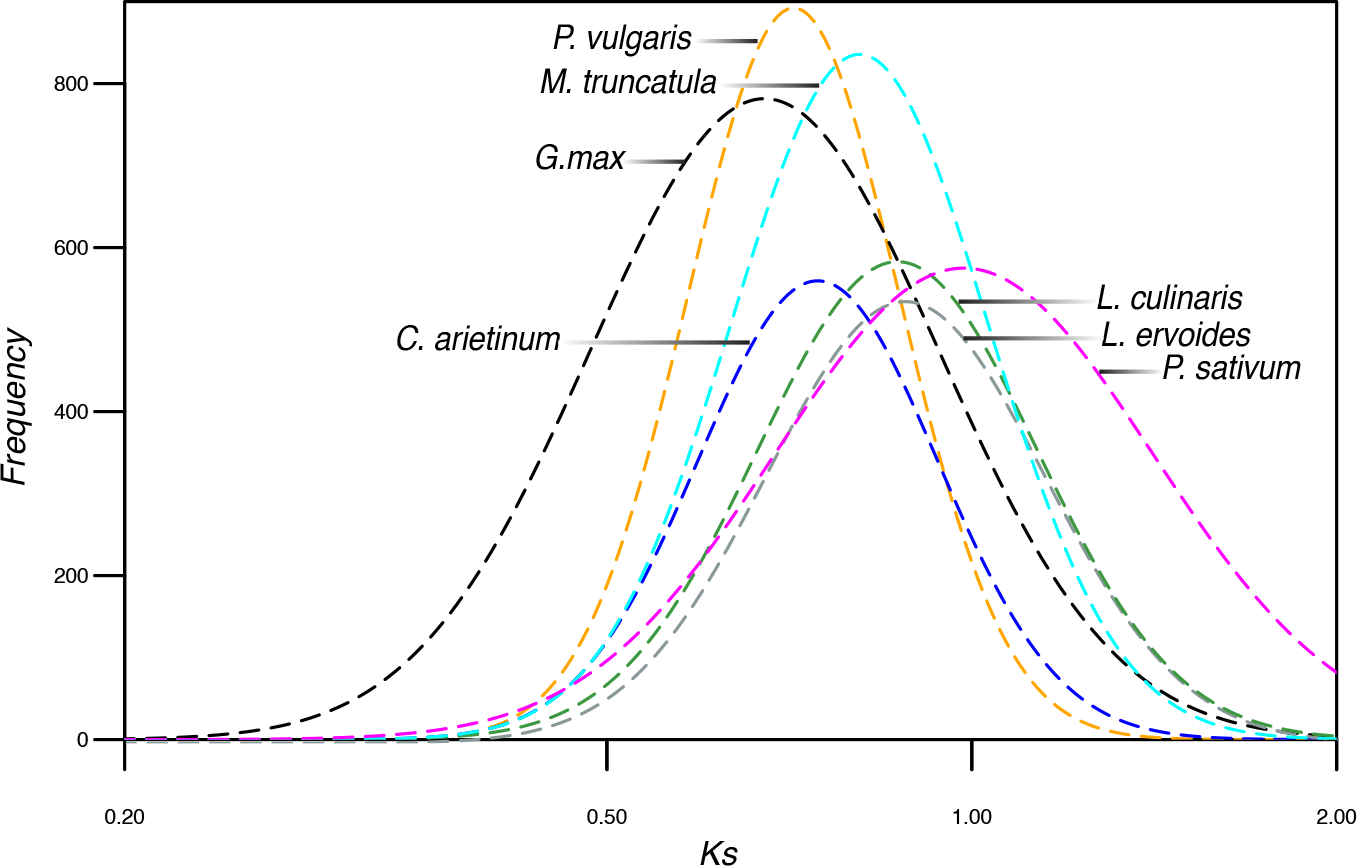
Mutation rate (Synonymous substitution rate) in legume species. The mutation rate was extrapolated based on the mean synonymous substitution (Ks) values of the WGD event shared by all legume species that occurred ∼56.5 Mya (Lavin et al., 2005. Systematic Biology 54: 575-594)

**Supp Fig 3.**
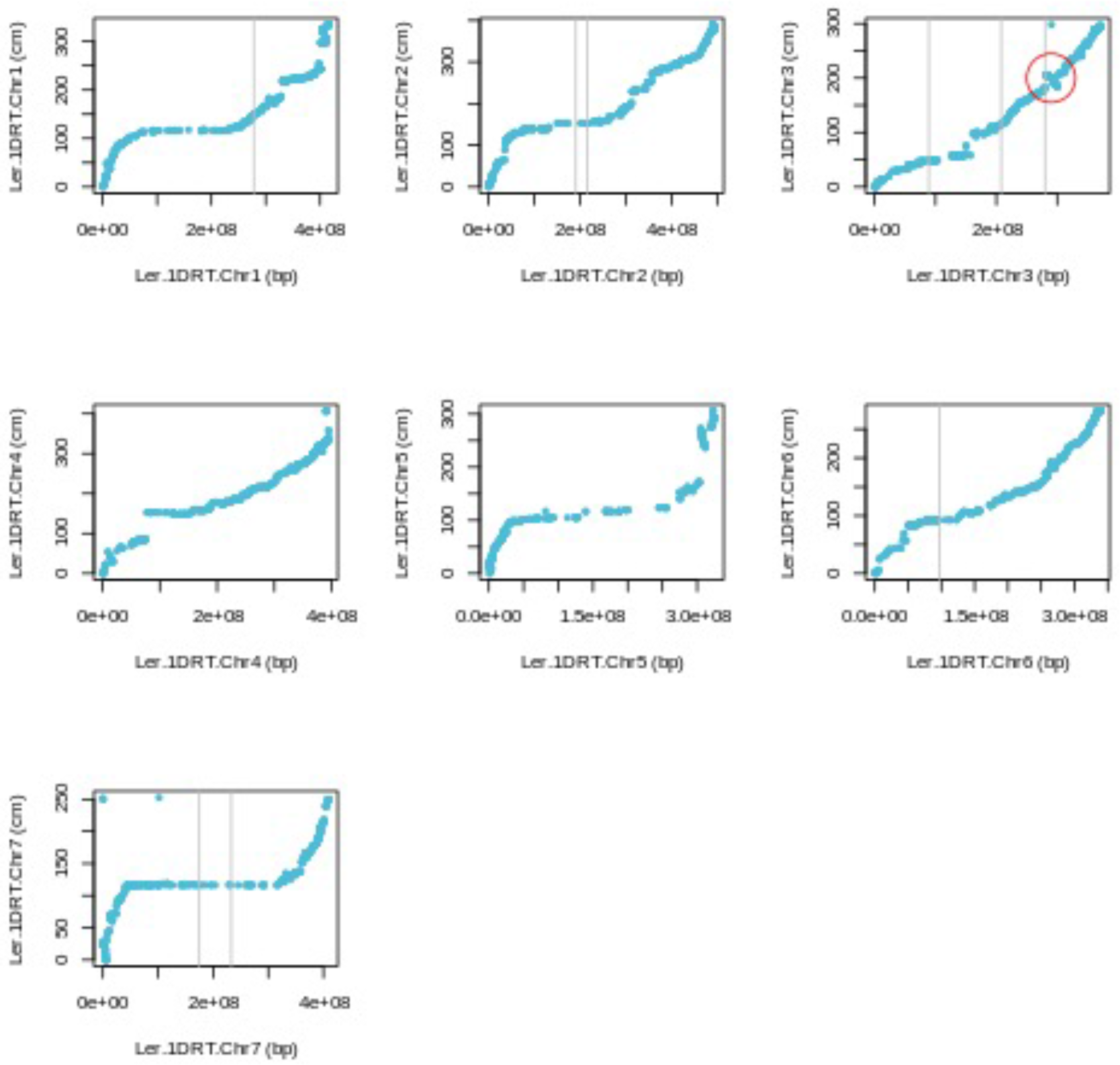
LR-66 linkage map marker cM versus Ler.1DRT genomic positions. Grey lines indicate the position of lentil-specific centromeric satellite repeats associated with centromeric chromatin. Inversion related to the segregation bias towards *Lens ervoides* on chromosome 3 circled in red.

**Supp Fig 4.**
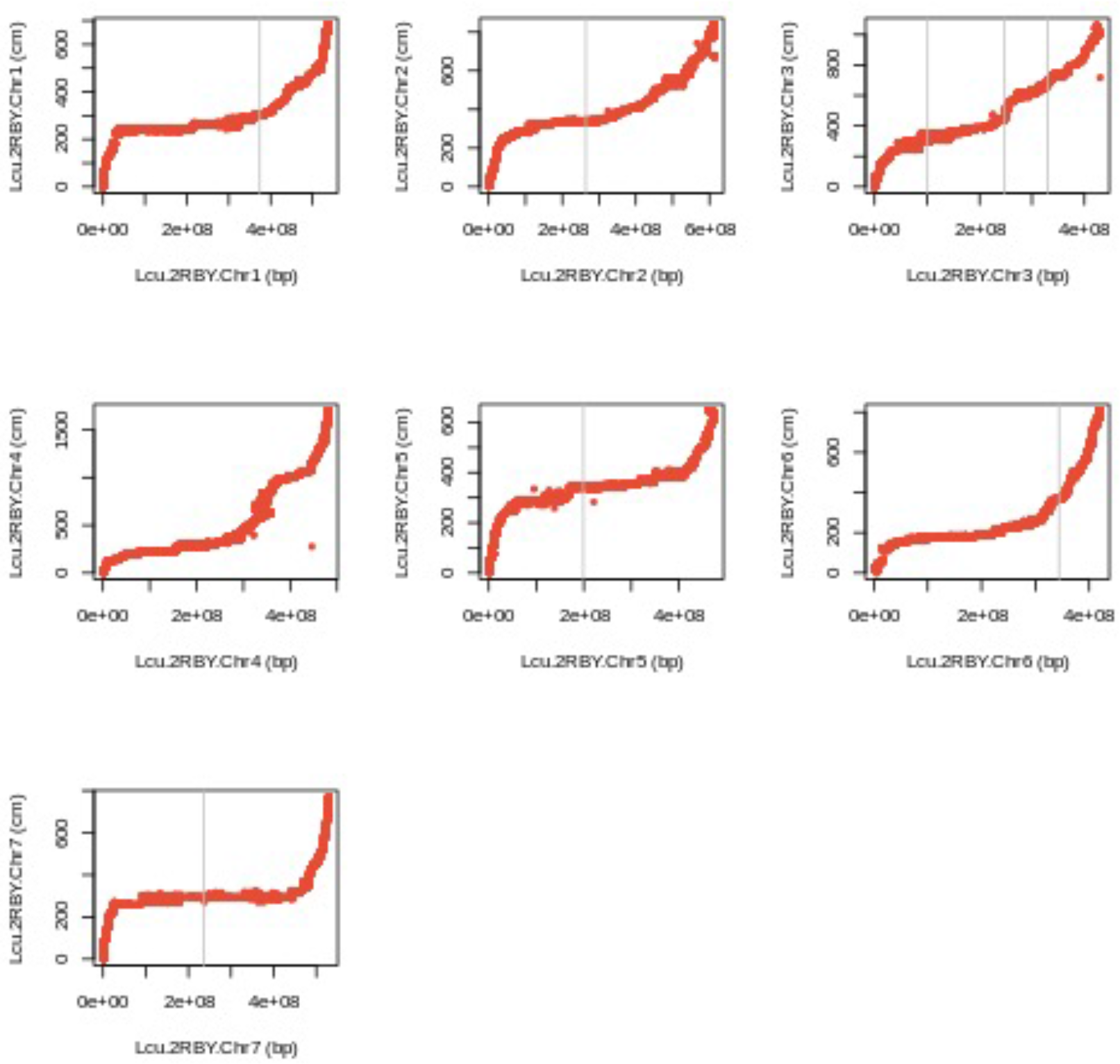
LR-01 linkage map marker cM versus Lcu.2RBY genomic positions. Grey lines indicate the position of lentil-specific centromeric satellite repeats associated with centromeric chromatin.

**Supp Fig 5.**
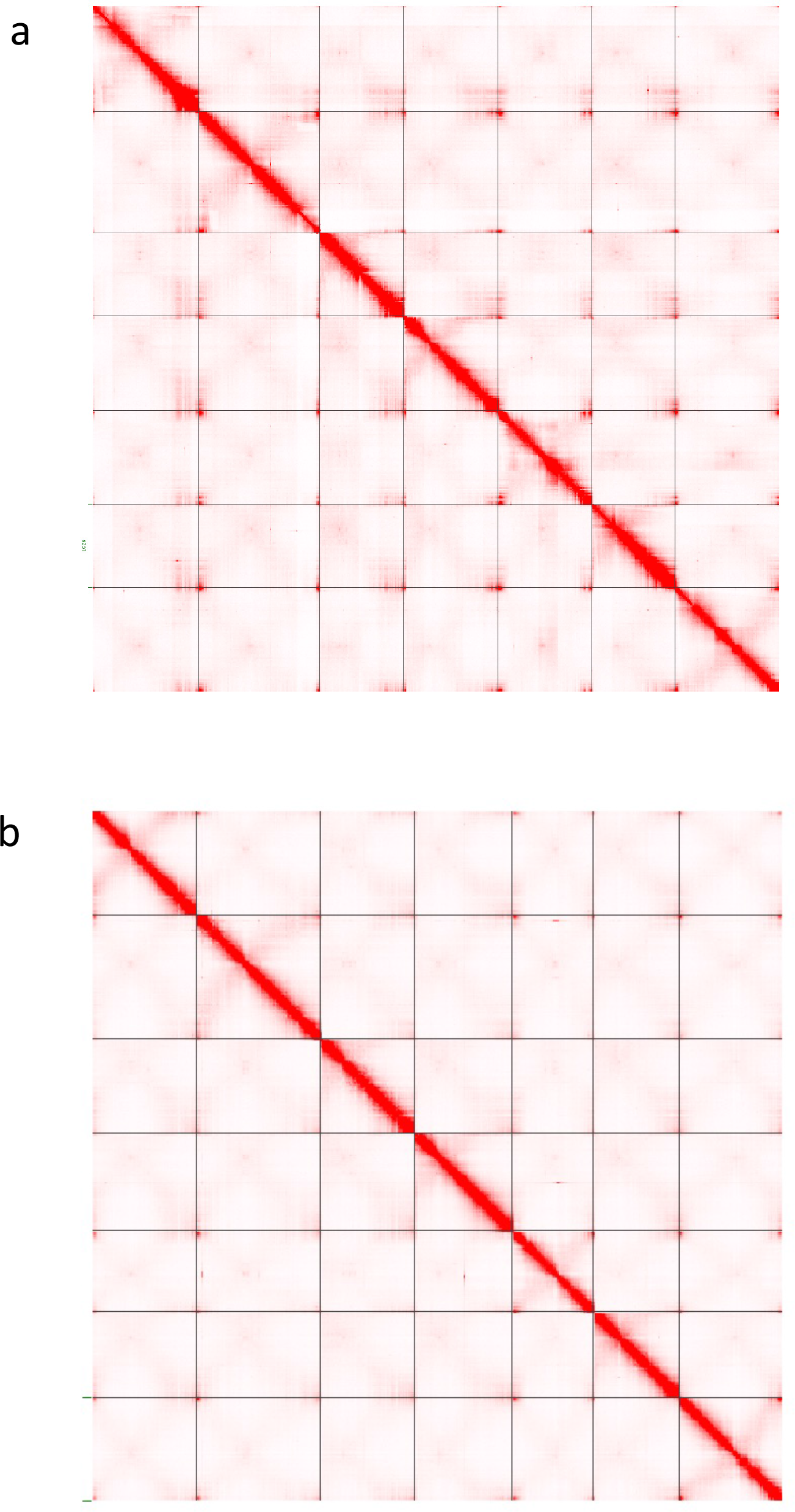
Hi-C contact maps for a) Lcu.2RBY and b) Ler.1DRT.

